# Embryonic stem cells do not have a globally disrupted higher-order chromatin fibre structure

**DOI:** 10.1101/2025.04.22.649930

**Authors:** James Ding, Shelagh Boyle, James Allan, Nick Gilbert

**Affiliations:** MRC Human Genetics Unit, Institute of Genetics and Cancer, The University of Edinburgh, Crewe Rd, Edinburgh, EH4 2XU, UK

**Keywords:** Chromatin, higher-order chromatin fibre, stem cell, pluripotency, biophysical, linker histones

## Abstract

Embryonic stem cells (ESCs) are thought to maintain pluripotency through global hyper-transcription, linked to an “open” chromatin structure. To investigate this idea, we analyzed higher-order chromatin fibres from mouse ESCs, their differentiated progenitors, and terminally differentiated mouse NIH3T3 fibroblasts. Bulk chromatin composition including protein:DNA ratio and nucleosome repeat length varied little between the cell types, but surprisingly biophysical analyses such as linker histone FRAP, hydrodynamic sedimentation and nuclease sensitivity also showed no significant differences in the conformation of purified higher-order chromatin fibres. To better evaluate the structure of higher-order chromatin fibres observed in cells, we developed a novel technique called SPOCC (Sedimentation Properties of Cross-Linked Chromatin). This approach revealed that ESCs and differentiated cells share similar bulk higher-order chromatin fibre structures, whilst ESCs have a slightly more disrupted structure than NIH3T3 cell chromatin. These results indicate that ESC transcriptional activity and plasticity are not driven by a fundamentally “open” higher-order chromatin conformation.

## Introduction

Since pluripotent mouse embryonic stem cells (ESCs) were first derived, they have been studied in detail and found to have many distinct characteristics^1^, including a short cell cycle^2^, sensitivity to DNA damage^3^, high levels of homologous recombination^4^, and low mutation rates^5^. In addition, stem cells express 40-60% of their genes, compared with only 10-20% of differentiated cells, and show high transcriptional heterogeneity, a property that is thought to provide them with pluripotent potential^6^. Stem cells are also highly transcriptionally active, in a state that is described as hypertranscription^7^ which might be important to drive the energetic demands of rapidly growing stem and progenitor cells during development. The basis for this is not fully understood, but it has been suggested that stem cells have a unique “open” chromatin fibre structure that is both plastic and dynamic and is necessary for pluripotency ^8^, and permissive for high levels of transcription^9–11^.

In eukaryotic cells, DNA is wrapped around histone proteins to form nucleosomes, which self-assemble into chromatin fibers^12^ primarily through electrostatic interactions^13^. The nucleosome building blocks are well described^14^, but the organisation of these components into higher-order structures are less well understood^15^, and in particular how they might influence development and differentiation. In vitro studies show that nucleosomes fold to form higher-order chromatin fibres that are approximately 30-nm in diameter^16,17^ with a helical configuration^18^. Structures analogous to this are observed in regions of the genome where nucleosomes are uniformly spaced, such as at centromeric satellites^19^, but across the majority of the genome and especially in genic regions, nucleosomes are irregularly positioned^20^, creating disruptions in the chromatin fibre giving it an irregular appearance^21^. Furthermore, the often-observed images of chromatin fibres belie their dynamic and flexible structure^22^ that is important to facilitate transcription factor binding, and engagement of the transcription machinery.

Chromatin is often characterised by the nucleosome repeat length, which is dependent on the amount of nucleosomes per length of DNA. Cell types that have a high transcriptional activity typically have more bound nucleosomes and a short repeat length, as observed for *S. cerevisae*^23^ or calf cortical neurons^24,25^. In contrast, ESCs have a repeat length of 186 bp^26^, which is reported to increase by 5-7 bp during cell differentiation^26^. Nucleosome positioning has also been characterised by mapping nucleosome dyads using chemical cleavage, which suggests that fragile nucleosomes can occupy previously identified nucleosome-depleted regions around transcription start sites of ES cells^27^. These promoter proximal nucleosomes are also often comprised of variant histones including H3.3 and H2AZ^28,29^ that bind to DNA with a reduced affinity^30^ and may create points of flexibility in the chromatin fibre^31^ that can facilitate access to chromatin complexes^32^, and endow the chromatin fibre with an ability to be more transcriptionally dynamic, mediated by the H2AZ acidic patch^33^.

The electrostatic folding of the chromatin fibre is dependent on linker histones^13,34^. Unlike core histones^35^, linker histones are highly dynamic^36,37^ with a globular domain that positions the globule domain of the protein near the dyad ^38^, and a long C-terminal tail that can shield charges along the fibre, facilitating folding. Linker histone levels affect chromatin folding: over-expression of linker histone H5 in rat sarcoma cells abrogates cell proliferation^39^ whilst a reduction in linker histones promotes global^40^ and local^41^ nuclear decompaction. Furthermore, H1 mutations are found in lymphoma and a reduction in linker histones show an activation of stem cell genes^42^, and C-terminal frame shift mutations in H1.4 leads to premature termination and gives rise to a neurodevelopmental disorder called Rahman syndrome^43^. The molecular mechanisms are not understood, but linker histone mutations or truncations could lead to altered protein binding affecting chromatin fibre structure and global levels of transcription. Consistent with this notion linker histone proteins have been shown to be highly dynamic in mESCs^9^, but with conflicting results^44^, and this rapid mobility is often interpreted to indicate that ESCs have a unique and “open” or “loose” higher-order chromatin fibre^8^; a feature which is proposed to contribute to their developmental plasticity^45,46^. Consistent with this notion GFP-H2B shows greater fluctuations in stem cells than differentiated cells, indicative of an altered nuclear architecture^47,48^. Early microarray studies also demonstrated that ESCs are transcriptionally hyperactive and this has been suggested to be a consequence of a “loose” chromatin organisation of ESCs^11^. Consistent with this observation, FAIRE-seq shows that stem cells have more disrupted or nucleosome free regions compared to stem cells^49^. At gross levels of nuclear organization stem cells do appear to undergo a change in architecture upon cell differentiation with a reorganization of chromocentres^9,50^ and a concomitant change in chromatin texture that has been suggested to be influenced by the Chd1 chromatin remodelling complex^46^. Furthermore, correlative electron spectroscopic imaging suggest that stem cells have more dispersed 10-nm like fibres both in euchromatic^51,52^ and heterochromatic regions^53^.

To explore the “open chromatin” hypothesis we examined bulk chromatin fibres isolated from stem cells and their differentiated progenitors using a combination of microscopic and biophysical approaches^12,51^. Contrary to expectation, we were unable to find any differences between bulk higher-order chromatin fibres analysed using biophysical sedimentation assays. Furthermore, we find that linker histones have a similar mobility in stem cells and differentiated cells. Our data suggests that the packaging of bulk higher-order chromatin fibres does not contribute to the hyper-transcription observed in ESCs or their unlimited potential to differentiate.

## Results

### Sucrose gradient sedimentation discriminates between chromatin fibres of different structures

To explore how chromatin architecture might contribute to the high levels of transcription observed in mESCs and to investigate the “open chromatin” hypothesis, we studied the structure and conformation of intact chromatin fibres isolated from stem cells and differentiated cells^8,9,46^. Sucrose gradient sedimentation, a widely used method for characterizing the biophysical properties of soluble chromatin fibers^31,54,55^, was used to compare and contrast chromatin fibre structures. In this approach, chromatin fibres are isolated from cell nuclei by briefly digesting the chromatin with micrococcal nuclease under physiological conditions, and gently releasing the soluble chromatin. After removal of nuclear debris, the chromatin is fractionated on a 6-40% sucrose gradient centrifuged for 3 h in an SW41 rotor (Fig 1A). Under these conditions the chromatin will sediment in the sucrose gradient, based on its mass and chromatin fibre structure. Large and therefore heavier fibres will sediment more rapidly, whilst smaller and light fibres will sediment more slowly. Importantly though, two fibres of the same mass and the same structure will sediment at the same rate, but if these fibres have different structures they will sediment at different rates. Essentially, this is because the sedimentation rate of an unfolded fibre be retarded due to its friction in the solvent, compared to a more compact fibre. Previously, we showed that mouse and human satellite containing chromatin fibres sedimented more rapidly than bulk chromatin, and by using simple modelling we argued that satellite containing chromatin adopted a rigid-rod like structure, whilst the bulk chromatin from the genome was interspersed by disruptions making it more flexible.

**Figure 1.**
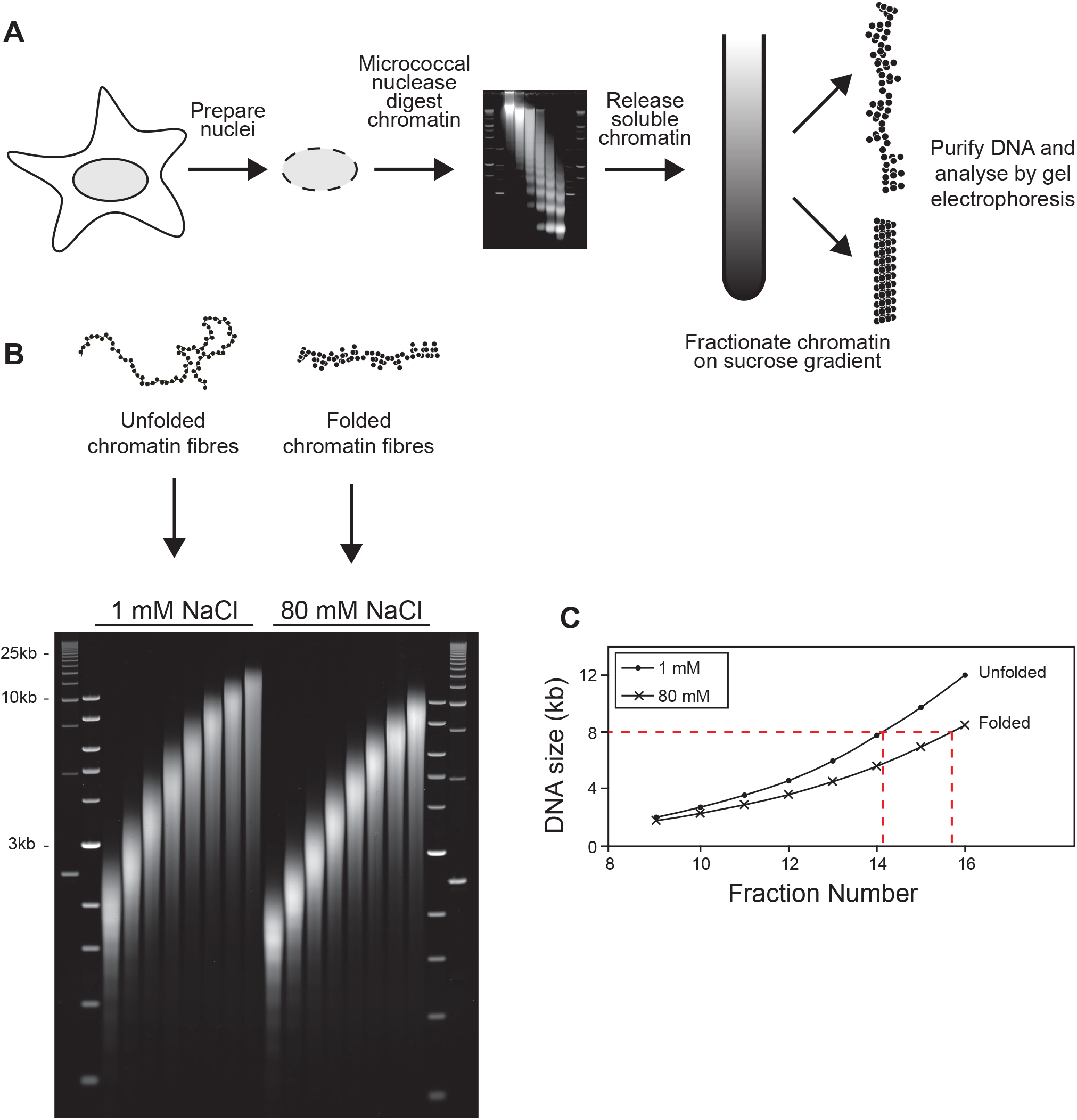
Differently folded higher-order chromatin fibres sediment at different rates. A. Schematic showing the isolation of soluble chromatin from NIH3T3 cells, and fractionation in a sucrose gradient in the presence of 1 mM NaCl or 80 mM NaCl. 80 mM NaCl is equivalent to physiological salt conditions, whilst chromatin fibres unfold in 1 mM salt. B. Chromatin prepared from NIH3T3 cells and fractionated in an isokinetic sucrose gradient in the same centrifuge in the presence of either 1 mM NaCl or 80mM NaCl. DNA was purified from individual fractions from each gradient and fractionated on an agarose gel, stained with ethidium bromide and scanned on a Fuji FLA-5000 laser scanner. C. Graph showing the relationship between DNA size and fraction number (sedimentation rate) for differently folded chromatin fibres. Red line shows how a 8 kb chromatin fibre with an unfolded (or disrupted) structure sediments more slowly (fraction ∼14), whilst a 8 kb chromatin fibre with a folded (or compact) structure sediments more rapidly (fraction ∼16).

To demonstrate that sucrose gradients can discriminate between chromatin fibres of different structures soluble chromatin was isolated from NIH3T3 cells, and dialysed either into a low salt (1 mM NaCl) or physiological salt (80 mM NaCl) buffer. Under these conditions chromatin in low salt is unable to shield the electrostatic interactions, causing the chromatin fibre to adopt an unfolded configuration^34,56^. After sedimentation the chromatin was fractionated, DNA was purified from individual fractions and analysed by agarose gel electrophoresis. DNA fragment size was plotted against fraction number (Figure S1C) enabling the relationship between the size and sedimentation rate to be determined. Consistent with expectations, for equivalent sized chromatin fibres (e.g. 6 kb) the sedimentation of the fibre unfolded in 1 mM NaCl was significantly retarded compared to when the fibre was folded in physiological salt, demonstrating that sucrose gradient sedimentation can discriminate between chromatin fibres of different structures.

### Stem cell and differentiated cell chromatin proteins

Sucrose gradient sedimentation can discriminate between chromatin fibres of different structure (Fig 1). As the sedimentation rate is affected by the mass of the chromatin fibre, we characterised the protein composition and density of chromatin. Total nuclear proteins were extracted from mouse ESCs, differentiated ESCs and mouse NIH3T3 embryonic fibroblasts and fractionated by SDS-PAGE (Fig 2A, left panel). Levels of core histones and linker histones were similar between the samples, so to examine proteins directly bound to the chromatin fibre, soluble chromatin was isolated from cells and fractionated on a sucrose step gradient (Fig S1A).

**Figure 2.**
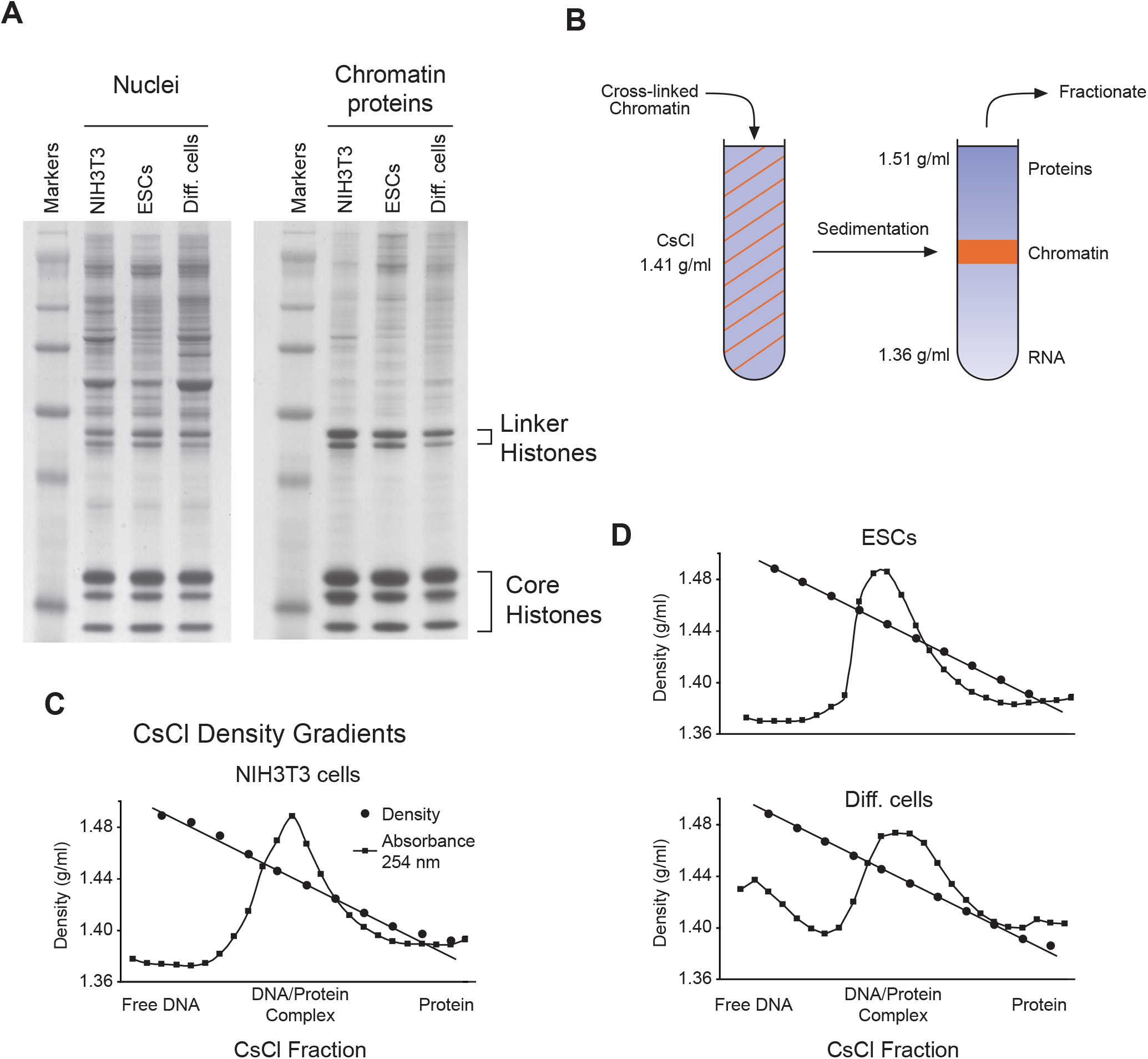
Protein and DNA composition of ESC and differentiated cell chromatin. A. Proteins were extracted from nuclei isolated from NIH3T3 cells, embryonic stem cells (ESCs) and differentiated ES cells. Chromatin proteins were extracted from chromatin that had been purified on a sucrose step gradient. Proteins were fractionated by 12% SDS-PAGE and stained with Coomassie Brilliant Blue. B. Cartoon depicting the sedimentation of chromatin in a caesium chloride gradient to determine chromatin density (i.e. protein:DNA ratio). C-D. Graphs showing the relationship between chromatin sedimentation in a caesium chloride gradient and fraction density for NIH3T3 cells, ESC and differentiated cell chromatin. The position of chromatin (DNA/protein complex) can be estimated from the DNA absorbance of each fraction. See also Figure S1.

To ensure that soluble chromatin fibres isolated from cells were representative of the entire genome chromatin was prepared from ESCs, differentiated ESCs and NIH3T3 cells. Soluble chromatin was purified on a 10-50% step gradient (Fig S1B) and the DNA purified (Fig S1C). When the DNA probes were hybridised to mouse metaphase spreads^55^ they gave a uniform staining pattern indicating that the population of fibres being studied are representative of the genome (Fig S1D-E). After fractionation of chromatin on a step gradient (Fig S1B), proteins were isolated from the peak fraction, concentrated by ethanol precipitation and analysed by SDS-PAGE. As for the nuclei, all histone bands were very distinct (Fig 2A, right panel), however, there were slight differences between the linker histone and core histone ratios between independent protein preps but as linker histones are very sensitive to degradation these differences might be a consequence of some protein degradation. To quantitatively analyse chromatin density (essentially the protein:DNA ratio), isolated chromatin fibres were purified on sucrose step gradients (Fig S1B), dialysed into triethanolamine buffer and extensively cross-linked using formaldehyde (Fig 2B). After cross-linking, samples were analysed by isopycnic centrifugation in caesium chloride gradients^54^. By measuring the refractive index and DNA absorbance of individual fractions the density of the purified chromatin fibres was calculated (Fig 2C-D). NIH3T3 cells and differentiated ESCs had very similar densities of 1.44 g/ml whilst stem cell chromatin was slightly denser at 1.45 g/ml. To put this into context this is equivalent to an increase in mass of the nucleosome by 0.7%, or one extra linker histone for every 12 nucleosomes.

### Characterising bulk nucleosome repeat length

Nucleosome position influences gene transcription by modulating accessibility to the underlying DNA, whilst high nucleosome density correlates to high transcriptional activity ^24,25^. Multiple factors affect nucleosome repeat length including linker histones^57^, and their loss promotes a reduction in repeat length and global chromatin decondensation leading to an increase in nuclear size^40^. Furthermore, short repeat length chromatin does not readily bind linker histone because the altered trajectories of linker DNA sterically impairs H1 binding^58^. For this reason, the repeat lengths of stem cells and their differentiated counterparts were compared using a nuclease digestion time course. As nucleosome repeat length is difficult to measure accurately, we have developed two new approaches to improve results. Historically micrococcal nuclease was used to digest chromatin to single nucleosomes^54,59^ but it has a tendency to nick within the nucleosome and trim the nucleosome ends. We therefore used DFF/CAD nuclease to digest chromatin^60^ (Figure 3A). DFF/CAD nuclease precisely cuts chromatin to yield nucleosomal oligomers and trims nucleosomes to a far lesser extent. After digesting nuclei, DNA was purified and fractionated by agarose gel electrophoresis (Fig 3B-C). The sizes of the nucleosomal bands were accurately measured for each time point and the data were plotted as band number against fragment size (Fig 3D-F). The gradient of this relationship provides the nucleosome repeat length for each time point of digestion. As a second improvement, nuclei were digested with different concentrations of enzyme enabling us to extrapolate back to calculate the nucleosome repeat length (Figure 3C-E). NIH3T3 cells, stem cells and differentiated cells all have very similar nucleosome repeat lengths of approximately 200-206 bp consistent with what we have observed previously using micrococcal nuclease^54^. In a previous study, Teif et al.^26^, calculated nucleosome repeat lengths from next generation sequencing data for MEFs, ESCs and differentiated ESCs as 191, 186 and 193 bp, respectively. This is consistent with our result, considering their experiment was undertaken with micrococcal nuclease which will have trimmed the DNA fragments to an extent. Our data therefore suggests that although transcription patterns change dramatically between ESCs and differentiated ESCs it is not accompanied by a substantive change in nucleosome repeat length, and by extension nucleosome density.

**Figure 3.**
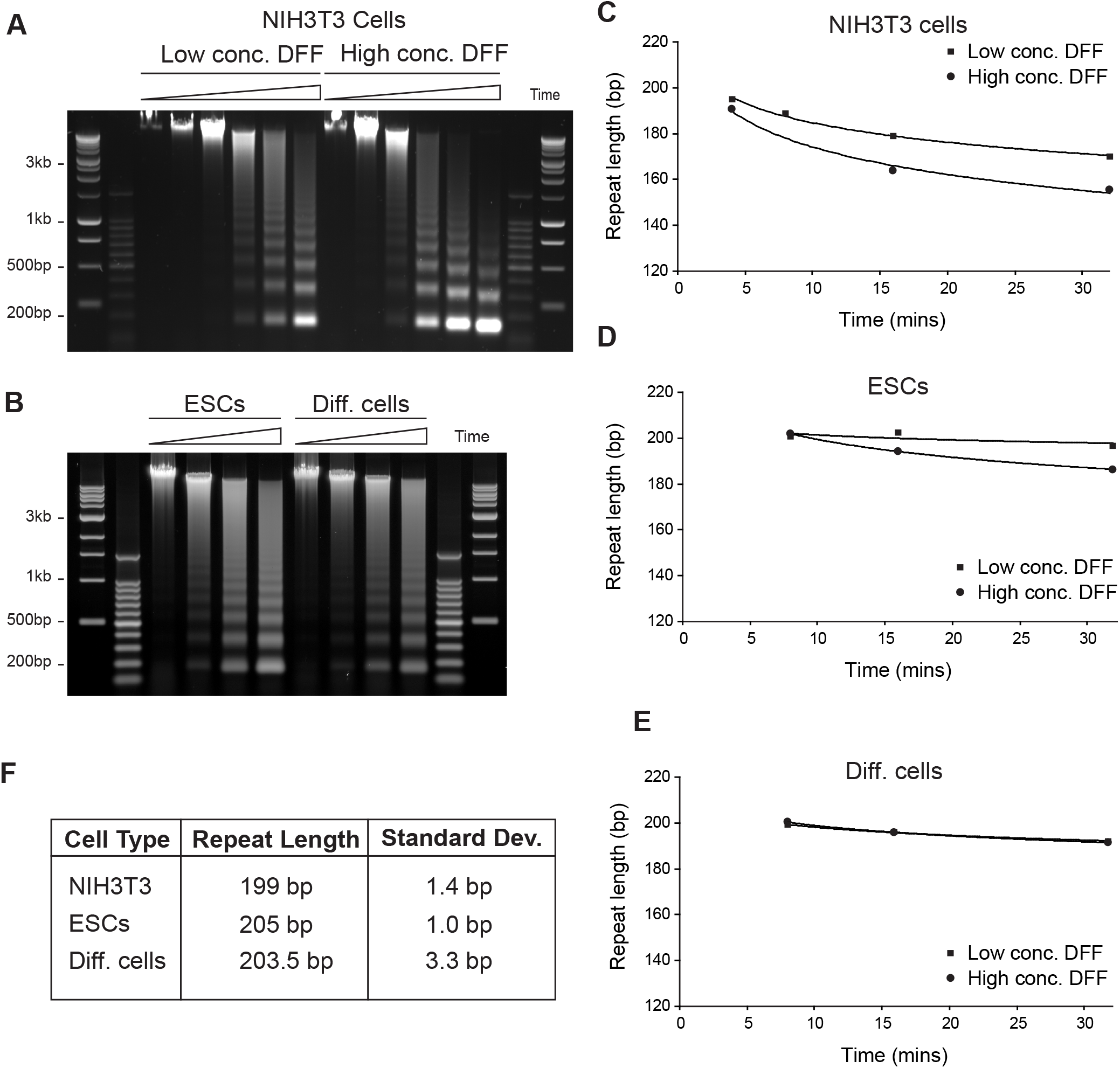
Nucleosome repeat length of ESC and differentiated cell chromatin. A-B. Nuclei samples digested with DFF nuclease, DNA digested and fractionated on an agarose gel. C-E. Graphs showing the nucleosome repeat length of chromatin digested for different amounts of time with different concentrations of DFF/CAD nuclease. F. Table showing a summary of the nucleosome repeat lengths of NIH3T3 cells, ESCs and differentiated cells.

### Hydrodynamic sedimentation of purified stem cell and differentiated cell chromatin

To establish whether soluble bulk higher-order chromatin fibres isolated from ESCs were more ‘open’ or ‘disrupted’ than chromatin from NIH3T3 fibroblasts, the hydrodynamic properties of soluble chromatin fibres isolated from these two cell types were compared. To achieve this, nuclei were digested briefly with micrococcal nuclease and large soluble chromatin fragments were released overnight. After clarification, the soluble chromatin was centrifuged in identical 6-40% isokinetic sucrose gradients in the same rotor in TEEP_80_ buffer, which will maintain chromatin in its native physiologically folded state (Figure 1). After sedimentation the chromatin was fractionated and DNA was purified from individual fractions and the DNA size analysed by agarose gel electrophoresis (Figure 4A). Plotting DNA fragment size against fraction number (Figure 4B) showed that ESC and NIH3T3 cell bulk chromatins have equivalent sedimentation rates suggesting that their levels of fibre compaction are very similar. Even for very large chromatin fragments of 14 kb, where it was anticipated differences in structure would be accentuated, no distinction between the two chromatin preparations were observed (Figure 4B).

**Figure 4.**
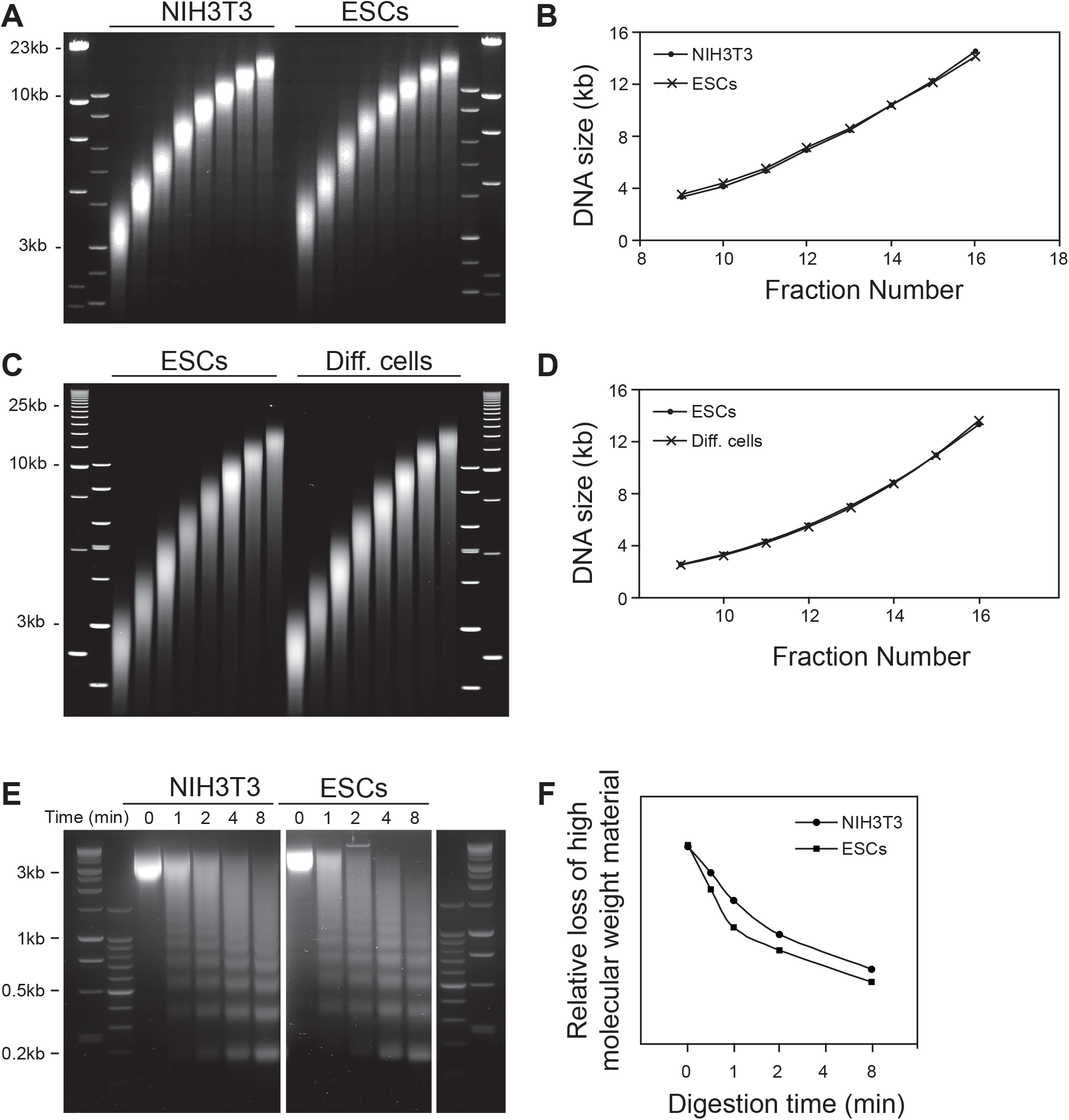
Chromatin fibres isolated from NIHT3 cells, stem cells and differentiated cells have similar hydrodynamic sedimentation properties. A. Chromatin was prepared from ES cells and NIH3T3 cells fractionated in an isokinetic sucrose gradient in the presence of 80mM NaCl. At this salt concentration the sucrose gradient will fractionate chromatin based on its mass and higher-order chromatin fibre structure. DNA purified from individual fractions from each gradient was analysed on an agarose gel, stained with ethidium bromide and imaged on a laser scanner. B. Graph showing the relationship between DNA size and fraction number (sedimentation rate) determined from gel shown in (A) for chromatin isolated and fractionated from NIH3T3 cells and ES cells, (D) or from ES cells and differentiated ES cells. C. As for (A) but samples were ES cells and differentiated ES cells. D. As for (B), however experiment compared samples from ES cells and differentiated ES cells. E. Soluble chromatin was isolated from NIH3T3 and ES cells and purified on a sucrose gradient. The isolated chromatin was digested with micrococcal nuclease over time and the DNA from individual samples was purified and fractionated on an agarose gel. The gel was stained with ethidium bromide and scanned on a laser scanner. F. The decrease in the signal intensity of the starting material was analysed and plotted against time for the two samples.

Although ESCs and NIH3T3 cells have very different differentiation potential they are both stable, highly proliferating cell lines. It is therefore possible that over a period of time in culture there has been a change in the conformation of the differentiated NIH3T3 cell chromatin to a “standard” higher-order chromatin fibre organisation similar to that found in ESCs. To test this idea, the conformation of chromatin fibres from ESCs was compared directly to cells derived from differentiated ESCs using retinoic acid. Soluble chromatin was isolated from the cells and fractionated on a sucrose gradient and the DNA sizes of individual fractions were compared (Figure 4C-D). Against expectations, chromatin fibres isolated from ESCs and differentiated cells sediment at similar rates, suggesting that have very similar structures.

### Nuclease sensitivity of chromatin isolated from stem cells and differentiated cells

A detailed hydrodynamic examination of bulk higher-order chromatin fibres isolated from undifferentiated and differentiated cells failed to reveal any differences in their chromatin fibre structure (Figure 4A-D). However, previous nuclease sensitivity studies have suggested that there is a difference in the chromatin compaction between stem cells and differentiated cells^46,61^. Our preliminary experiments indicated that nuclease digestion of different nuclei preparations gave unreproducible results due to the variable permeability of different cell types to the enzymes. To circumvent this challenge gradient purified chromatin was used as the substrate for nuclease digestion.

Chromatin prepared from NIH3T3 cells and ESCs was fractionated through a sucrose gradient. Equivalently sized fractions were taken from each gradient and digested with a micrococcal nuclease time course (Figure 4E). The digestion rate was determined by quantifying the loss of high molecular weight DNA (> 1 kb) over time (Figure 4F). Results show that ESC chromatin is digested slightly faster than NIH3T3 cell chromatin, indicating that there might be small differences in the chromatin fibre structure.

### Linker histone binding in stem cells and differentiated cells

Linker histone binding to nucleosomes shields electrostatic interactions to compact higher-order chromatin fibres in vivo^13,34,62,63^. This binding predominantly occurs between the linker histone globular domain^38,64^ and the nucleosome dyad, which is abolished by mutating key residues^65^. Previously we used fluorescence recovery after photobleaching (FRAP) of linker histone H1 and H5 to investigate their interactions with native and unmethylated chromatin fibres^59^. We showed that linker histones were more stably bound to chromatin in the absence of DNA methylation, however, we were unable to find a concomitant alteration in higher-order chromatin structures. Nevertheless, a reduction in nuclear linker histone levels promotes global chromatin decondensation and an increase in nuclear size^40^ whilst localised H1 binding compacts chromatin and represses gene expression^41^. As Fig 4F showed differences in nuclease sensitivity we wondered if this might be dependent on linker histone binding. It has also been reported that linker histone binding to stem cell chromatin is hyperdynamic and it has been proposed that this property is a barrier to establishing higher-order chromatin structures and contributes to the maintenance of stem cell plasticity^9^. We therefore used FRAP to investigate linker histone binding in ESCs and differentiated ESCs cells. Linker histones H1.4 and H5 were tagged with GFP at their N termini, adjacent to an alpha helix that interacts with linker DNA at the entry/exit point of the nucleosome^38^ and minimising any interference of GFP with the high-affinity C-terminal chromatin binding domain of H1^66,67^ (Figure 5A). Although H1 and H5 are similar, H5 is more highly charged and binds to chromatin with a higher affinity^68^. Because of this H5 may better discriminate between different extents of chromatin binding than H1 (Figure 5A). ESCs were transfected with GFP-H1, GFP-H5 or a binding mutant of GFP-H5 (Figure 5B) and linker histone mobility was analysed by following fluorescence recovery every 7 s after photobleaching (3 sec) (Figure 5C). The recovery curves and kinetics of H1 (t_1/2_ = 30 sec) and H5 (t_1/2_ = 42 sec) in ESCs (Figure 3D) are similar to what we observed previously^59^. By FRAP, H1 mobility is identical in ESCs and differentiated ESCs. In contrast the H5 binding mutant is exceeding mobile (t_1/2_ = 6 sec). Histone H5 appeared to be more mobile (t_1/2_ = 33 sec) in differentiated stem cells rather than ESCs (t_1/2_ = 42 sec), but as the initial rates of recovery (2.7 - 2.9 relative fluorescence units per sec) were very similar between cells types, these results indicate that linker histone mobility did not significantly vary between stem cells and differentiated cells, suggesting that differences in linker histone binding would be unlikely to be a major determinant in higher-order chromatin fibre compaction.

**Figure 5.**
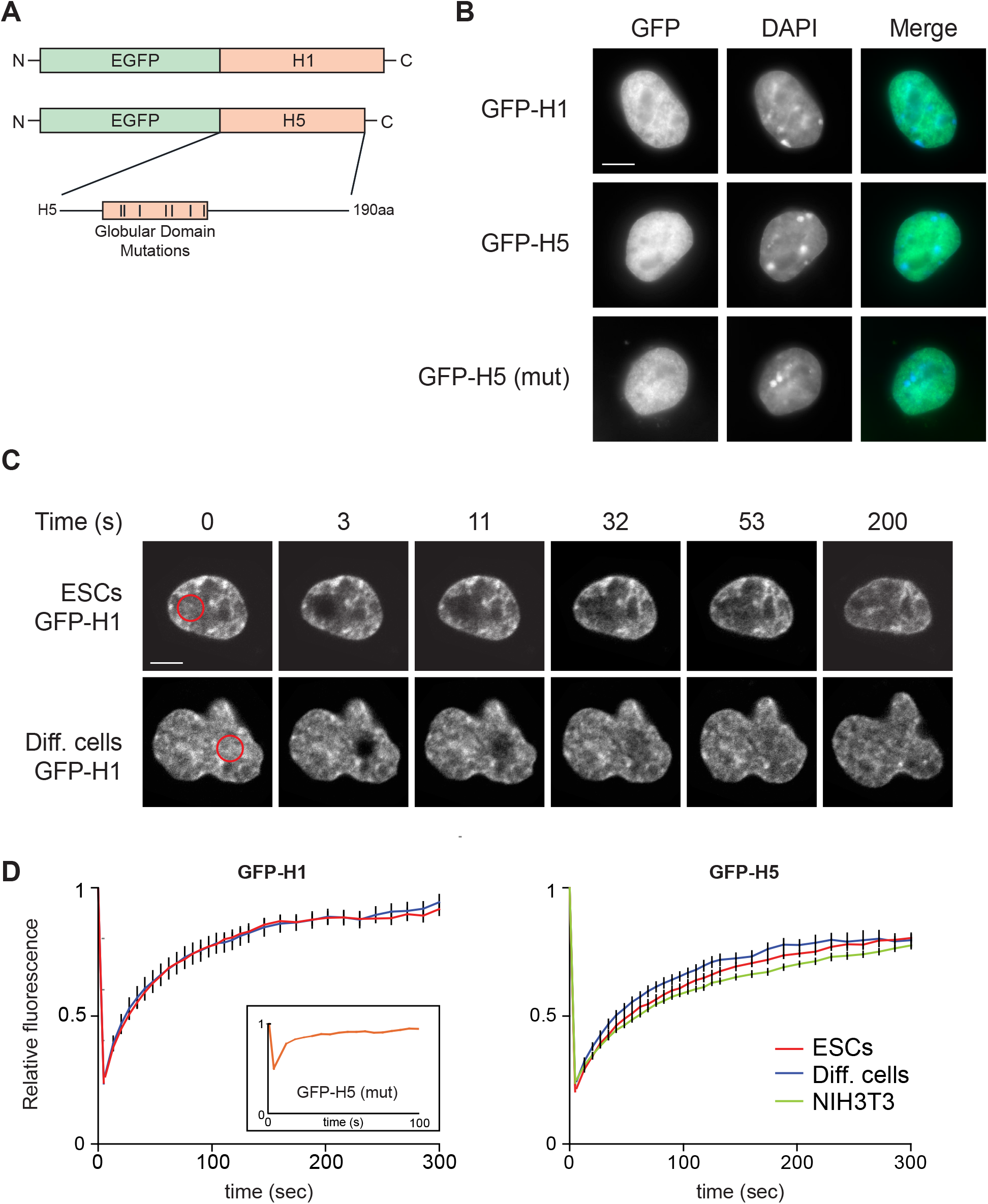
Linker histone mobility in ESCs and differentiated cells. A. Schematic showing linker histones H1 and H5 tagged at their N-termini with EGFP and the position of the globular domain H5 mutations. B. Distribution of GFP fluorescence (green) in ES cells transfected with GFP-H1, GFP-H5, or GFP-H5 mutant. DNA is counterstained with DAPI (blue). Scale bar, 5 μm. C. Representative confocal images of GFP-H1 transfected ES cells or differentiated ES cells during FRAP. The bleach area is marked by red circles and the fluorescence recovery is monitored over time. D. Relative fluorescence within the bleach ROI during FRAP of GFP-H1 (left), GFP-H5 (right) or GFP-H5 mutant (inset) expressed in ES cells (red), differentiated ES cells (blue) or NIH3T3 cells (green). Graphs show mean values (±SEM, error bars). 3T3 H5, n = 21; ES H1, n = 32; Diff H1, n= 43; ES H5, n= 32; Diff H5, n = 41.

### Characterisation of in vivo fixed stem cell and differentiated cell chromatin

The studies described so far have compared the structure of unfixed chromatin fibres isolated from NIH3T3 cells, ESCs and differentiated cells (Figure 4) using sucrose gradient sedimentation. While evidence suggests that these buffer conditions closely mimic the nuclear environment ^34,55^ and accurately preserve the structure of higher-order chromatin fibres, there remains a possibility that isolating chromatin fibres alters their conformation, potentially obscuring structural differences. To address this limitation, we developed an approach to investigate chromatin fibres cross-linked within cells to preserve their native structure, we refer to this method as SPOCC (Sedimentation Properties of Cross-Linked Chromatin).

To evaluate SPOCC we undertook a series of control experiments. Firstly, formaldehyde cross-linking was evaluated to ensure it would maintain the structure of the higher-order chromatin fibre (Fig S2A). Chromatin fibres were prepared from NIH3T3 cells, dialysed into either low salt (1 mM) or high salt (80 mM) conditions to alter their structure. These fibres were extensively cross-linked with 1% formaldehyde, then dialysed into a buffer containing 1 mM NaCl/tris to quench and remove any residual formaldehyde. In the absence of cross-linking low-salt conditions would be expected to unfold chromatin fibres. After sedimentation chromatin fibres were isolated by fractionation, cross-links reversed, DNA purified and examined by agarose gel electrophoresis (Fig S2B). By plotting the relationship between DNA size and fraction number (Fig S2C) we showed that despite sedimentation in low salt buffer, chromatin fibres cross-linked in 80 mM NaCl sedimented more rapidly (compare 6 kb fibres in Fig S3C; 1 mM fibres sediment in fraction 13, 80 mM fibres sediment in fraction 16). This result demonstrates that extensive formaldehyde cross-linking effectively preserves distinct chromatin fibre structures.

Secondly, formaldehyde cross-linking generates extensive intra- and inter-chromatin fibre cross-links, substantially reducing the ability to release chromatin fibres from cells. To address this, we hypothesized that digesting histone tails with trypsin after extensive cross-linking could facilitate chromatin fibre release. Soluble chromatin was isolated from NIH3T3 cells, dialyzed into low- and high-salt buffers, and extensively cross-linked using formaldehyde. The chromatin was then dialyzed to remove formaldehyde and lightly digested with trypsin (Fig S3A), resulting in the clipping of histone tails (Fig S3B). Furthermore, cross-linking was similarly efficient on unfolded or folded chromatin fibres based on the appearance of new protein bands in the 1 mM and 80 mM samples, whilst the protease digested the histones to their trypsin resistant core^69^. Chromatin fibres were solubilised in SDS and fractionated on sucrose gradients prepared in 1 mM salt. After sedimentation, gradients were fractionated (Fig S3C), cross-links reversed, DNA purified, and analysed by agarose gel electrophoresis (Fig S3D). Analysis of the sedimentation rates for the chromatin fibres (Fig S4E) indicated that the combination of cross-linking and subsequent trypsin digestion maintains detectable differences in chromatin fibre structure.

As our pilot experiments suggested that a combination of cross-linking and trypsin digestion did maintain the structural characteristics of chromatin, we decided to use SPOCC methodology to characterise the properties of cellular chromatin. To explore the reproducibility of the protocol two samples of NIH3T3 cells were analysed in parallel. Nuclei were digested with micrococcal nuclease and then extensively cross-linked using formaldehyde. After quenching with tris and dialysis to remove residual cross-linker, the samples were digested with trypsin, soluble chromatin was released in the presence of SDS and fractionated on a sucrose gradient. Fractions were collected, DNA purified and analysed by agarose gel electrophoresis (Fig S4A). The distribution of the DNA sizes from pairs of fractions were compared (Fig S4B) and the peak DNA size was determined for each fraction. After plotting fraction (sedimentation rate) against peak DNA size (Fig S4C) the two samples, as expected, were found to have very similar sedimentation properties, showing the reproducibility of the method for two independent samples.

SPOCC (Sedimentation Properties of Cross-Linked Chromatin) was then used to compare ESC and NIH3T3 cell chromatin. Cross-linked and trypsin digested chromatin was prepared from ESCs and NIH3T3 cells, and the histone proteins examined by SDS-PAGE to ensure they were similarly digested (Fig 6B). The soluble chromatin was fractionated based on its size and structure on a sucrose gradient and DNA from individual fractions analysed by agarose gel electrophoresis (Fig 6C). The distribution of DNA sizes from individual fractions were compared (Fig 6D-E) and indicated that ESC chromatin was marginally more disrupted than NIH3T3 chromatin, consistent with nuclease sensitivity observed in Fig 4F. These results suggest that SPOCC can discriminate between different chromatin structures, but also shows that some aspects of chromatin folding are lost if the samples are not cross-linked before analysis (Fig 4A). It is well known that some regions of the genome are more heterochromatic than other regions that are transcriptionally active^70^. Similarly, polycomb enriched regions are known to have characterisristics of heterochromatin and be potentially more difficult to extract from cells. To ensure that the chromatin released for analysis was representative of the genome purified soluble chromatin was fragmented using MNase and analysed by genome-wide next generation sequencing. The chromatin released was evenly distributed across the genome (Fig S5A). To further characterise the distribution of released chromtin we used the chromHMM classification (Fig S5B). Compared to genomic DNA isolated from B-cells there was enrichment in more disrupted portions of the genome such as enhancers and transcribed regions, but there was no obvious depletion in specific regions of the genome that might confound subsequent analysis.

**Figure 6.**
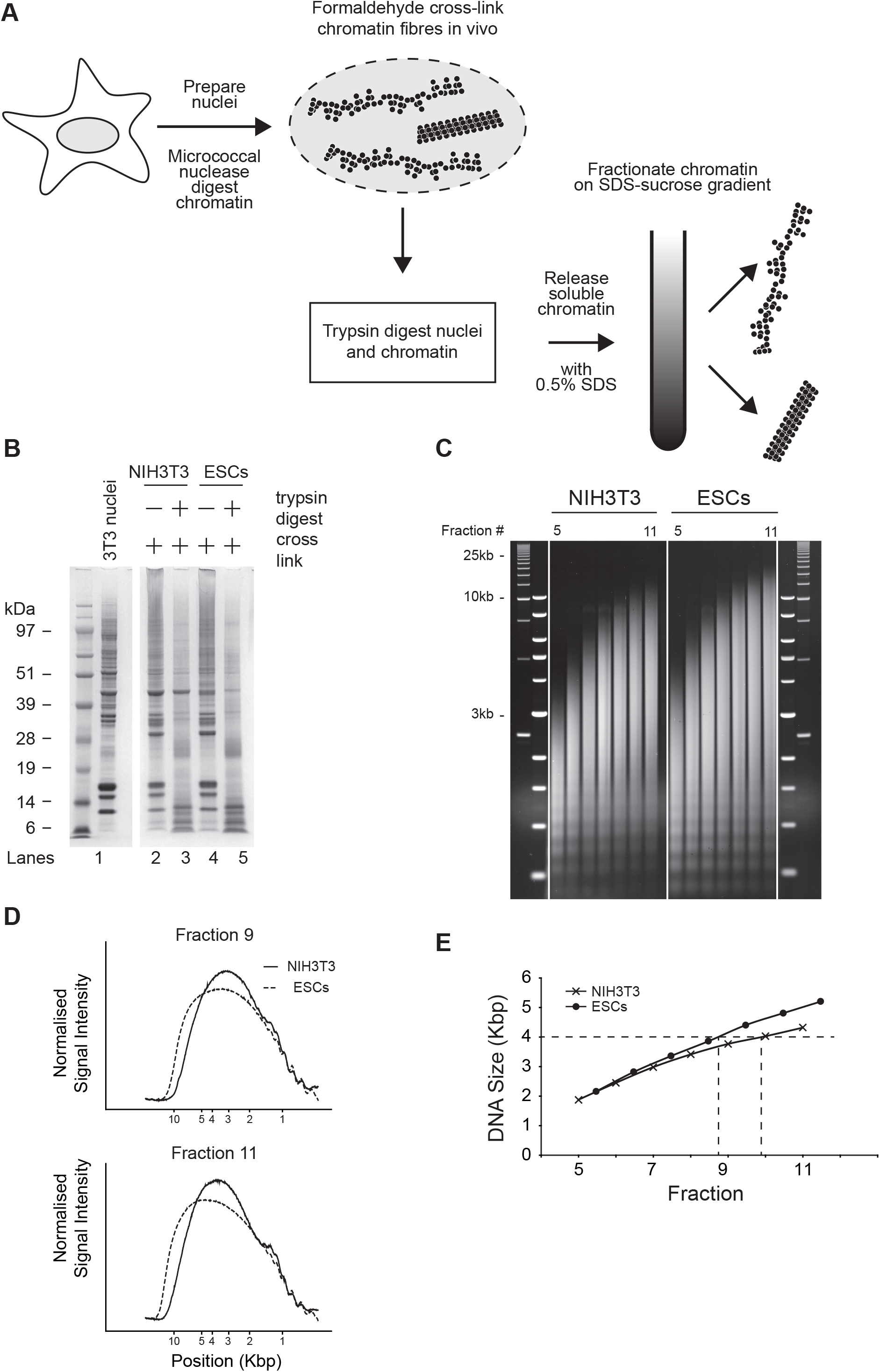
Sedimentation characteristics of chromatin fibres isolated from NIH3T3 and ES cells. A. Schematic showing the preparation, isolation and characterisation of chromatin fibres cross-linked inside cells. Nuclei are digested with MNase and then cross-linked extensively with formaldehyde to maintain intra-fibre cross-links. To isolate the chromatin inter-fibre cross-links are broken by digestion with trypsin and the chromatin is solubilised and fractionated based on its (fixed) structure in a sucrose gradient. B. Qualitative analysis of nuclei proteins isolated from cross-linked NIH3T3 and ES cells in the presence or absence of trypsin. Gel was stained with Coomassie blue. C. DNA isolated from individual sucrose gradient fractions analysed on an agarose gel, stained with ethidium bromide and imaged on a laser scanner. D. Densitometry of fraction 14 (top) and fraction 16 (bottom) for NIH3T3 and ESCs DNAs. E. Graph showing the relationship between DNA size and fraction (sedimentation rate), determined from the gel shown in (C). See also Figure S2-S5.

As ESCs and NIH3T3 cells are very different cell types, we decided to directly compare stem cells and differentiated stem cells, using our SPOCC method. Nuclei were prepared from ESCs and differentiated cells, digested with micrococcal nuclease and cross-linked. The chromatin was solubilised and fractionated on a sucrose gradient (Fig 7A), subsequently DNA was isolated from individual fractions and analysed on an agarose gel (Fig 7B). Analysis of individual fractions (Fig 7C) and for all fractions indicated that chromatin fibres from stem cells and differentiated cells sediment at the same rate, indicating that they have very similar bulk chromatin fibre structures.

**Figure 7.**
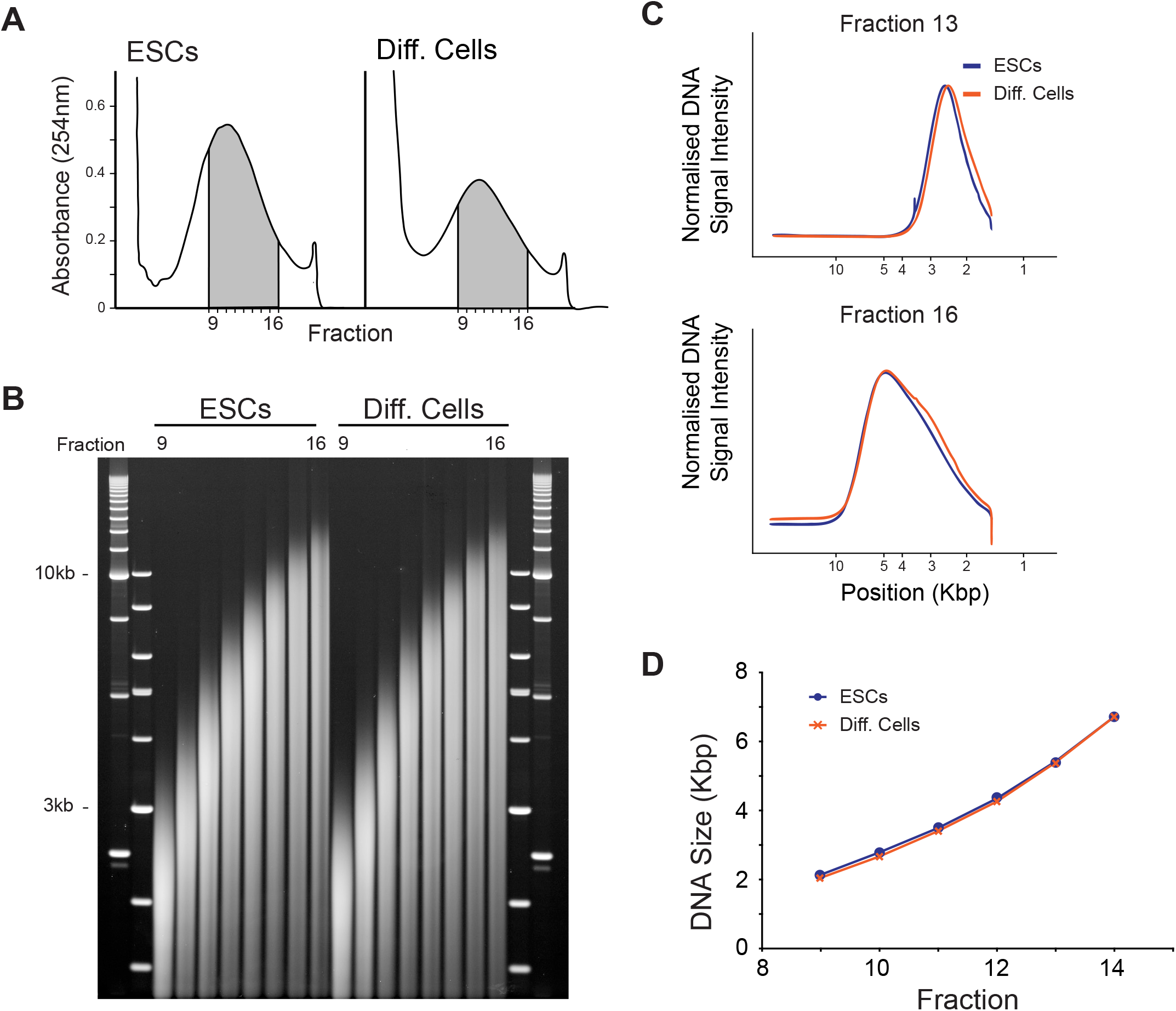
Sedimentation properties of stem cell and differentiated cell chromatin fibres cross-linked inside nuclei. A. Cross-linked chromatin isolated from ES cells and differentiated cells and fractionated in an isokinetic sucrose gradient in the same centrifuge in the presence of 1 mM NaCl. Graphs showing the absorbance (254 nm) across the sucrose gradients (top of gradient is left side). B. DNA purified from individual fractions from each gradient (grey area in A) was fractionated on an agarose gel, stained with ethidium bromide and scanned on a laser scanner. C. Densitometry of ethidium bromide staining in individual lanes shown in panel C. D. Relationship between DNA size and sedimentation rate (fraction) for stem cells and differentiated cell chromatin.

## Discussion

Our study indicates that bulk chromatin fibres isolated from stem cells and differentiated stem cells have equivalent structures (Fig 7D), contrary to the accepted dogma, and instead suggesting that the organization of the higher-order chromatin fibre does not contribute to the pluripotent stem cell state, *per se*. Synthetic disrupted chromatin fibres form ‘puddles’^17^, that by super-resolution imaging in cells appear to look like ‘clutches’^71^. Pluripotent cells have less dense clutches containing fewer nucleosomes than differentiated cells, suggesting that this difference might be linked to stem cell state. However, the hydrodynamic sedimentation approaches used in this study failed to detect differences in the conformation of the higher-order chromatin fibre between stem cell and differentiated cell bulk chromatin. This is unlikely to be due to a limitation in the technique as this approach has the ability to detect small differences in chromatin fibre folding in the vicinity of the promoter in transcriptionally active genes^31^, and we were able to observe that ESCs had a slightly more disrupted bulk chromatin fibre structure than NIH3T3 cells (Fig 6). Although these differences are difficult to quantify they are indicative of additional disruptions in the chromatin fibre. Furthermore, when chromatin was analysed in conditions favourable to higher-order folding, satellite-containing heterochromatin has a sedimentation rate approximately 20% greater than of bulk chromatin^54^. In this study we were unable to find any substantive evidence for differences in bulk chromatin between stem cells and differentiated cells, therefore, it may only be in specific regions of the genome where there are differences in the conformation of the higher-order chromatin fibre, such as in satellite-containing centromeric heterochromatin^54,72,73^, or in localised regions around transcription start sites^31^, or at transcriptionally active genes^74^. In these situations nucleosomes are dynamically remodelled to facilitate access to transcription factor binding sites and docking of the RNA polymerase. These regions are nuclease sensitive and readily degraded by DNaseI, revealing regions, that are often referred to as hypersensitive sites. Mapping studies suggest that most active genes have regions of disrupted chromatin covering approximately 150 – 200 bp. These regions are unlikely to be detected in this study for two reasons. Firstly, they are too small to have a significant enough effect on the sedimentation of chromatin fibres. Secondly, because these regions are sensitive to nucleases they are likely to be degraded by MNase early on in the digestion. Although chromatin remodeling machines like Chd1 play a role in maintaining stem cell genes in a special “open” state^10,46^, remodelling enzymes are unlikely to have a global effect on chromatin structure. Our results here are consistent with stem cell and differentiated cells having broadly similar levels of chromatin fibre compaction.

Previously, we estimated the extent of differences in chromatin fibre structure between bulk chromatin, satellite containing chromatin^54^, and chromatin observed at promoters^31^. It would similarly, be interesting to determine the molecular basis for difference in bulk chromatin structure between NIH3T3 cells and ESCs (Fig 6). To do this however it would be necessary to map different chromatin structures at higher resolution, using for example, next generation sequencing. This would indicate whether the differences observed in the experiments here are limited to specific genomic regions or are more global. Although the basis for this difference is not known, one possibility is that as ESCs replicate more rapidly than NIH3T3 cells the chromatin does not fold into such a regular structure. It is well known that H1.0 is expressed in terminally differentiated cells^75^, compacting chromatin^76^ and repressing transcription^77^. Recent reports suggest that fibroblasts express increased amounts of H1.0 which might have the effect of compacting the chromatin fibre^76^.

Human embryonic stem cells and iPS cells have great therapeutic potential to provide a source of specific cell types for the treatment of diseases. Central to the use of stem cells is our ability to understand the cellular processes required to regulate their differentiation pathways. This study demonstrates that although stem cells are poised for differentiation along different lineages their bulk higher-order chromatin fibre is a structure predominantly used for the regular packaging of DNA, rather than providing a generally permissive environment for gene expression. Instead, we suggest that it is the binding of master transcription factors to key regulatory genes that are responsible for stemness, and are sufficient to impose or re-impose a stem cell state without the need to modify global higher-order chromatin fibre structures.

## Limitations of the study

Sucrose gradient sedimentation has been used by us^31,54,55,59,78^ and other researchers^79^ to evaluate the structural properties of soluble chromatin fibres. In this approach, soluble chromatin is isolated from cells and then characterised outside of its cellular context. Although this allows questions to be asked of the fundamental principles of chromatin folding, there is a significant possibility that it does not reflect all aspects of chromatin folding inside the nucleus. Not least, many chromatin proteins are lost when the chromatin is isolated, supercoiling is also released when the chromatin is cut by nucleases, and nuclear processes such as transcription are significantly abrogated. To overcome these limitations for studying cellular chromatin, it would be better if chromatin could be “fixed” within the confines of the nucleus before analysis. Chromatin fixation is straightforward, but isolation of fixed chromatin is challenging. In this study we developed SPOCC, a technique which enables chromatin fibres fixed in the nucleus to be isolated and examined. To achieve this, cross-linked nuclei were briefly digested with trypsin to remove histone tails, and solubilised using SDS. Although, we provide controls to demonstrate that this method can still discriminate between chromatin fibres with a different structure, there is a possibility that other aspects of chromatin folding will also be lost. Not least, fixation and manipulation of the chromatin fibre will affect chromatin fibre dynamics, similarly, fixation might work with different efficiency in different parts of the nucleus. It is therefore important to interpret results using SPOCC with an understanding of its potential limitations and complement it where possible with other experimental techniques.

## Resource Availability

### Lead contact

Requests for further information and resources should be directed to and will be fulfilled by the lead contact, Nick Gilbert.

### Material availability

- This study did not generate new unique reagents.

### Data and code availability

- All data reported in this paper will be shared by the lead contact upon request.
- This paper does not report original code
- Any additional information required to reanalyze the data reported in this paper is available from the lead contact upon request.

## Acknowledgements

The authors would like to thank members of our research groups for useful discussions and for Dirk-Jan Kleinjan and Catherine Naughton for commenting on the manuscript. We also acknowledge Matt Pearson of the Advanced Imaging Resource at the Institute of Genetics and Cancer for his technical support. This work was supported by the UK Medical Research Council (MC_UU_00035/6).

## Author contributions

Conceptualization, N.G. and J.A.; methodology, J.D. and N.G.; Investigation, J.D., S.B. and N.G.; writing – original draft, N.G. and J.A.; writing – review & editing, J.D., S.B. and N.G.; funding acquisition, N.G.

## Supplemental information

## STAR Methods

### Key resources table

| REAGENT or RESOURCE | SOURCE | IDENTIFIER |
| --- | --- | --- |
| Antibodies |  |  |
| Rhodamine anti-dig sheep antibody | Vector |  |
| Texas Red anti-sheep antibody | Vector |  |
| Biotinylated anti-avidin antibody | Vector | BA-0300 |
| Chemicals, peptides, and recombinant proteins |  |  |
| DFF/CAD nuclease | Synthesised in lab |  |
| Mouse C<sub>0</sub>T-1 DNA | Roche |  |
| Avidin-FITC | Vector |  |
| Trypsin | Sigma | T8128 |
| Soya bean trypsin inhibitor | Sigma | T9003 |
| Experimental models: Cell lines |  |  |
| NIH3T3 cells |  |  |
| OS25 mouse embryonic stem cells | Billon et al., 2002 |  |
| ht2 mouse embryonic stem cells | Austin Smith (University of Exeter) |  |
| Recombinant DNA |  |  |
| DFF45 and DFF40 subunits cloned in pRSFDuet | Xiao et al., 2007 |  |
| GFP-H1 | Gilbert et al., 2007 |  |
| GFP-H5 | Gilbert et al., 2007 |  |
| GFP-H5 mutant | This study |  |
| Software and algorithms |  |  |
| IPLab |  |  |
| Bowtie2 v2.5.3 |  |  |
| Bedtools v2.31.1 |  |  |
| ChromHMM | https://compbio.mit.edu/ChromHMM/ |  |

### Experimental model and study participant details

The cell lines used in this study were NIH3T3 mouse embryonic fibroblasts, ht2 (*Oct-hyg-tk*) and OS25 ESCs (*Oct-hyg-tk*, *Sox2βgeo*)^80,81,84^.

### Method Details

#### Cell culture and differentiation

NIH3T3 cells were grown in high-glucose DMEM supplemented with 10% FCS and 1% pen/strep in 5% CO_2_. ESCs were grown on plastic coated with 0.1% gelatin in GMEM supplemented with 10% FCS, 2 mM glutamine, 100 mM sodium pyruvate, 1% pen/strep, 1% non-essential amino acids, 100 µM 2-ME and 100U/ml LIF in 5% CO_2_. OS25 ESCs were differentiated with retinoic acid (RA) using an abbreviated 5-day protocol^80,82^. For this protocol 5×10^−6^ cells were plated in a 100 mm dish in complete media and grown overnight. On day 1 media was changed with LIF being replaced by 5 µM RA, and on day 3 fresh media supplemented with RA was added. Cells were harvested on day 5. Gene expression was followed using RT-PCR and expression arrays^83^.

#### Nuclei and chromatin preparation and nuclease digestion

Nuclei and chromatin were prepared as described previously^54,55^. To determine nucleosome repeat length nuclei were prepared and diluted to 4 A_260_ units/ml in NBR buffer and were digested with 10-40 units/ml Mnase or 10-40 μl/ml DFF/CAD nuclease^60^. Aliquots were removed into stop buffer at suitable times and DNA was purified and analysed on 1.2% agarose gels in TBE buffer. To prepare soluble chromatin nuclei were diluted to 20 A_260_ units/ml in NBR buffer and digested with 6-10 units/ml MNase. Digestion was stopped with 10 mM EDTA and the nuclei were resuspended in TEEP20N buffer (10 mM Tris-HCl pH 8.0, 1 mM EDTA, 1 mM EGTA, 250 μM PMSF, 20mM NaCl, 0.05% NP40) overnight. For chromatin purification nuclear debris was removed by 5 min centrifugation in a microfuge and 850 μl soluble chromatin was layered on a 10% / 50% sucrose step gradient in TEEP80 buffer (10 mM Tris-HCl pH 8.0, 10 mM EDTA, 1 mM EGTA, 250 μM PMSF, 80mM NaCl) and centrifuged at 4°C (50,000 rpm) for 1hr 50 min in a MLS-50 rotor in a benchtop ultracentrifuge (Beckman). For chromatin structure analysis nuclear debris was removed by centrifugation and 400 μl soluble chromatin was loaded on to a 6-40% isokinetic sucrose gradient and centrifuged at 4°C (41,000 rpm) for 3.5hrs in a SW41 rotor in TEEP80 buffer. 500 μl fractions were collected from the gradients by upward displacement. DNA was purified from chromatin by phenol-chloroform extraction whilst proteins were purified by ethanol precipitation or extraction with Tri-Reagent (Sigma) followed by ethanol precipitation. Proteins were analysed on 12% NuPAGE gels (Invitrogen)

#### Caesium chloride density centrifugation

Soluble chromatin was purified on a 10%/50% step gradient as described above. The peak chromatin fraction (500 μl) was equilibrated into TEAP80 buffer (10 mM Triethanolamine-HCl pH 8, 1 mM EDTA, 80 mM NaCl, PMSF) using a MiniTrap Sephadex G-25 spin column (Cytiva), supplemented with 0.5% formaldehyde and rotated overnight at 4°C. The density of the samples were adjusted to 1.42 g/ml by adding 0.8 vol. saturated caesium chloride (1.92 g/ml) and centrifuged in a Sorval TV-865 rotor at 20°C (55,000 rpm) for at least 40 hrs. 250 μl fractions were collected from the bottom of the isopycnic gradients, refractive index (RI) was measured using a refractometer (Bellingham Stanley) and the DNA concentration was determined by measuring A_260_. Caesium chloride density = 10.819 × RI – 13.441.

### Histone H1 and H5 constructs

Plasmids for GFP-H1 and GFP-H5 were described previously^59^. To make a GFP-H5 binding mutant positively charged lysine and arginine amino acids in the H5 globular domain were replaced with either alanine or glutamic acid (K40E, R42E, K52A, K69A, R73A, K85A, R94A)^65^, the globular domain was then cloned into full length H5, and fused to GFP.

#### FRAP

ESCs and differentiated daughter cells were grown on 35 mm glass bottom dishes (Iwaki) and transfected with either GFP-H1, GFP-H5 or a GFP-H5 binding mutant. 24 h after transfection, samples were mounted onto a Nikon A1R, equipped with a Solent Scientific incubation chamber incorporating temperature and humidified CO_2_ control. The microscope comprised of a Nikon Eclipse TiE inverted microscope with Perfect Focus System and was equipped with a 457/488/514nm Multiline Argon lasers. Data were acquired using NIS Elements AR software (Nikon Instruments Europe, Netherlands).

Cells expressing high levels of H1- or H5-GFP fusion protein (total cellular pixel intensity > 35,000) were excluded from analysis. For FRAP, a 4-μm-diameter region of interest (ROI) of the nucleus in the midfocal plane was bleached for 3 s using the 488 laser (30%) power. Images were captured with a 60× objective at 7-s intervals for a total of 120 s and then 14 s intervals for 180 s using 10% of laser power. Each image was processed by an interactive script (IPLAB version 3.6; Scanalytics) to correct for nuclear rotation and cell movement. Loss of fluorescence attributed to the imaging process alone was assessed from the sum of pixel intensities in the cell. The fluorescence intensity for each ROI over time was then normalized to this.

#### FISH

To examine the representative release of soluble chromatin (Fig S2D-E), DNA purified from chromatin fractions was labelled with either biotin or digoxigenin and hybridised to 3:1 fixed metaphase spreads prepared from NHI3T3 cells. Mouse C_0_T-1 DNA was included in hybridisations to suppress signal from repetitive DNA. Biotinylated probes were detected using sequential layers of avidin-FITC and biotinylated anti-avidin. Digoxigenin-labeled probes were detected using rhodamine anti-dig sheep antibody and Texas Red-anti-sheep antibody (Vector). Slides were counterstained with 0.5μg/ml DAPI Slides were examined with an epifluorescence microscope (Axioskop; Carl Zeiss MicroImaging, Inc.) equipped with a 100× NA 1.3 lens and a CCD camera (Micromax; Princeton Instruments).

#### SPOCC

To examine the Sedimentation Properties of Cross-Linked Chromatin (SPOCC), nuclei were prepared and micrococcal nuclease digested as described above. Nuclei were washed in TEAP80 buffer and cross-linked in 1% formaldehyde 10 min at RT. The sample was boosted to give 2% formaldehyde and cross-linked at 4°C for 4 hrs on a wheel. Samples were dialysed against TEN20P (10 mM Tris-HCl pH 7.5, 1 mM EDTA, 20 mM NaCl, PMSF) overnight and digested with 40 μg/ml trypsin (Sigma T8128, 1500 units/mg). The reaction was stopped by adding 40 μg/ml soyabean trypsin inhibitor (Sigma T9003), PMSF and SDS to 0.5%. Samples were incubated 10 min at room temperature and then loaded on to a 6-40% isokinetic sucrose gradient and centrifuged at 4°C (41,000 rpm) for 4.5hrs in a SW41 rotor in TEEP1 buffer (10 mM Tris-HCl pH 8.0, 10 mM EDTA, 1 mM EGTA, 250 μM PMSF, 1mM NaCl). 500μl fractions were collected from the gradients by upward displacement and the DNA from them was purified by phenol-chloroform extraction and analysed by gel electrophoresis.

#### In vitro chromatin cross-linking and trypsin digestion

Soluble chromatin was prepared as described above and dialysed against TEAP1 (10 mM Triethanolamine-HCl pH 8, 1 mM EDTA, 1 mM NaCl, PMSF) or TEAP80 buffer. Chromatin in 1 mM NaCl adopts an unfolded, 10-nm, configuration whilst in 80 mM NaCl it is folded to give a 30-nm fibre^34^. To cross-link the samples formaldehyde was added to 1% and incubated at 4°C for 6 hrs on a wheel. Samples were then processed as for *in vivo* chromatin cross-linking and digestion.

#### DFF/CAD nuclease preparation

Engineered mouse DFF45 and DFF40 subunits were cloned in pRSFDuet (Novagen) (gift from W.Garrard, University of Texas Southwestern Medical Centre)^60^ to enable co-expression. The plasmid was transformed into BL21(DE3)-RP cells (Stratagene). Cells were grown until 2OD and then induced using 1 mM IPTG at 16°C. After 12 hours cells were harvested and lysed in Binding buffer (300 mM NaCl, 15 mM Imidazole, 50 mM Tris-HCl (pH 8), 10% Glycerol, 10 mM 2ME, PMSF, Protease inhibitor tablet). Lysozyme was added to 1 mg/ml for 30 min on ice and the cells were extensively sonicated. The mixture was clarified by adding triton-X100 to 0.1% and cleared by centrifugation. The protein was then purified on nickel agarose using standard techniques, desalted into storage buffer (100 mM KCl, 20 mM Tris-HCl pH 8.0, 0.2 mM EDTA, 2 mM DTT, 10% Glycerol) and diluted to give 50% glycerol. Protein quality was estimated by gel electrophoresis and activity was characterised by digesting naked DNA and nuclei. The DFF/CAD nuclease was activated by digestion with TEV enzyme (Invitrogen) at 30°C and was used in NBR buffer (5.5% sucrose, 85 mM KCl, 10 mM Tris-HCl pH 7.5, 1.5 mM CaCl_2_, 3 mM MgCl_2_, PMSF).

### Nuclease digestion of purified chromatin

A single isokinetic fraction of soluble chromatin purified by sedimentation in a 6-40% isokinetic sucrose gradient was supplemented with 3mM CaCl_2_ and digested with micrococcal nuclease (0.2 units/μg) at room temperature. An 80μl aliquot was removed into MNase stop buffer (1% SDS, 100μg/ml proteinase K, 2.5mM EDTA) for time 0. To the remaining material, micrococcal nuclease was added and 80μl aliquots were removed on a time course basis into MNase stop buffer. The DNA was purified for agarose gel electrophoresis. All of the chromatin samples tested negative for endogenous nucleases or residual micrococcal nuclease within the time-frame of these experiments.

### Agarose gel electrophoresis and analysis

Large DNA fragments were fractionated by gel electrophoresis in 0.7-1% agarose in 1× TPE buffer (90mM Tris-phosphate, 2mM EDTA) with buffer circulation whilst small DNA fragments were fractionated by gel electrophoresis in 1%-1.2% agarose in 1× TBE buffer (50mM Tris-borate, 1mM EDTA). Ethidium bromide stained agarose gels were scanned using a 532 nm laser and a 580nm band-pass filter on a Fuji FLA-5000. The size of the bands was determined from the DNA size markers which were a 2.5 kb DNA ladder (Bio-Rad) or a 1 kb or 100 bp DNA ladder (NEB) using Aida v3.22 analysis software.

### Analysis of chromatin genome-wide coverage

Soluble chromatin from mouse ES cells and differentiated cells was prepared as described above. DNA was extracted using SDS, proteinase K digestion, phenol:chloroform:isoamyl-alcohol extraction, and ethanol precipitation. DNA was electrophoresed on a 0.7% TPE gel for 24 Vh/cm and then extracted from a gel slice using beta agarase (NEB), focussing on 2-2.5 kb modal molecular weight DNA. This was used to generate libraries using the NEBNext® Ultra™ II FS DNA Library Prep, with minimal amplification (8 cycles). Sequencing was performed on illumina NextSeq P3 flow cell (Novogene), generating 50 bp paired end reads.

Alignment of reads to GRcm38 was performed using Bowtie2 (v2.5.3, –very-sensitive-local), followed by filtering to remove alignments with a MAPQ > 20. The mean depth of properly paired primary alignments was calculated for each 1 kb window of GRCm38 using bedtools coverage (v2.31.1). This was then normalised based on the number of filtered alignments, read length, and effective genome size (depth per genomic coverage, DPGC). Smoothing was performed, per chromosome, using a 1 Mb rolling median. A similar approach was taken to calculate the DPGC in chromatin states generated using E14 ChIP-seq data^85^.

For comparison PCR free whole genome sequencing data from three mouse strains^86^ was analysed using the same approach. Down sampling was performed to retain a random selection of 10% of reads.

## Quantification and statistical analysis

Statistical tests are detailed in figure legends. Analysis was performed in base R or Excel

## Additional Resources

None

## Supplementary figures

**Figure S1 (related to Figure 2)**

**Soluble chromatin fibres released from somatic cells, stems cells and differentiated stem cells are representative of the entire genome.**

A. Soluble chromatin was isolated from cells and purified on a 10%/50% sucrose step gradient. Fractions were collected by upward displacement.

B. Representative graph showing the absorbance (254 nm) of chromatin across a sucrose gradient (top of gradient is left side).

C. DNA was purified from individual gradient fractions and analysed by gel electrophoresis and stained with ethidium bromide.

D. Peak DNA fractions were taken and labelled by nick translation. Top, DNA isolated from soluble chromatin (green) was comparatively hybridised on a mouse metaphase spread with total genomic DNA (red) from the same cells in the presence of Cot-1. Bottom, enlarged images of selected chromosomes. Chromosomes were counterstained with DAPI (blue).

E. DNA isolated from ES cell soluble chromatin (red) was comparatively hybridised to a mouse metaphase spread with DNA isolated from differentiated ES cell soluble chromatin (green) in the presence of Cot-1. Chromosomes were counterstained with DAPI (blue).

**Figure S2 (related to Figure 6)**

**Differently folded and cross-linked chromatin fibres exhibit a pronounced difference in sedimentation rate.**

A. Schematic showing the experimental approach. Soluble chromatin fibres were isolated from NIH3T3 cells and either folded in 80 mM NaCl or unfolded in 1 mM NaCl. Chromatin structure was fixed by extensive formaldehyde cross-linking introducing intramolecular bonds. After cross-linking chromatin samples were fractionated on a sucrose gradient in low salt (1 mM NaCl).

B. Chromatin isolated from NIH3T3 cells, structure altered by changing salt concentration, fixed and fractionated in an isokinetic sucrose gradient. DNA was purified from individual fractions from each gradient, cross-links reversed, DNA purified and fractionated on an agarose gel, stained with ethidium bromide and scanned on a Fuji FLA-5000 laser scanner.

C. Graph showing the relationship between DNA size and fraction number (sedimentation rate) for differently folded and cross-linked chromatin fibres.

**Figure S3 (related to Figure 6). Differently folded, cross-linked, and trypsin digested NIH3T3 chromatin fibres have different sedimentation rates.**

A. Schematic showing the isolation of soluble chromatin fibres from cells. Chromatin fibres were either folded in 80 mM NaCl or unfolded in 1 mM NaCl. Chromatin structure was fixed by extensive formaldehyde cross-linking introducing intramolecular bonds. After cross-linking cross-linker was remove by dialysis, trypsin digested to remove tail-to-tail interactions and solubilised in SDS. Samples were then fractionated on a sucrose gradient in low salt (1 mM NaCl).

B. Coomassie stained protein gel showing nuclear proteins in total NIH3T3 nuclei (lane 1), chromatin proteins before cross-linking (lanes 2 and 3), after cross-linking (lanes 4 and 5) and after trypsin digestion (lanes 6 and 7).

C. Cross-linked chromatin isolated from NIH3T3 cells, and folded in either low (1 mM; unfolded) or high salt (80 mM NaCl; folded) and fractionated in an isokinetic sucrose gradient in the same centrifuge in buffer containing 1 mM NaCl. Graphs showing the absorbance (254 nm) across the sucrose gradients (top of gradient is left side).

D. DNA purified from individual fractions from each gradient (grey area in C) was fractionated on an agarose gel, stained with ethidium bromide and scanned on a laser scanner. Left, samples digested with high levels of trypsin, Right, samples digested with low levels of trypsin.

E. Densitometry of ethidium bromide staining for individual lanes shown in panel D to identify the size of peak fractions. Graph shows relationship between DNA size and fraction (sedimentation rate).

**Figure S4 (related to Figure 6)**

**Reproducibility of extracting cross-linked, trypsin digested NIH3T3 chromatin samples**

A. Cross-linked and trypsin digested chromatin was isolated from independent samples of NIH3T3 cells and fractionated on sucrose gradients in low (1 mM) NaCl. After fractionation DNA was purified from individual samples and fractionated on an agarose gel, stained with EtBr and quantified on a laser scanner (FLA-5100).

B. Densitometry of DNA intestity in individual lanes from (top) fraction 14 and bottom (fraction 16.

C. Graph showing the relationship between DNA size and fraction number for the sedimentation of independent NIH3T3 chromatin samples.

**Figure S5 (related to Figure 6)**

**Representative release of soluble chromatin from cells**

A. Soluble chromatin was purified from ES cells and differentiated ES cells and sequencec. Graph showing the distribution of released chromatin (in depth per genomic coverage) across each chromosome in the mouse genome, compared to the distribution of genomic DNA isolated from B cells from different mouse strains. Data shown is an average of two biological replicates, smoothed across 1 Mb windows.

B. Characterisation of chromatin into different genomic states using the chromHMM classification. Data shown is an average of two biological replicates.

