## Supplementary figures for "Embryonic stem cells do not have a globally disrupted higher-order chromatin fibre structure"

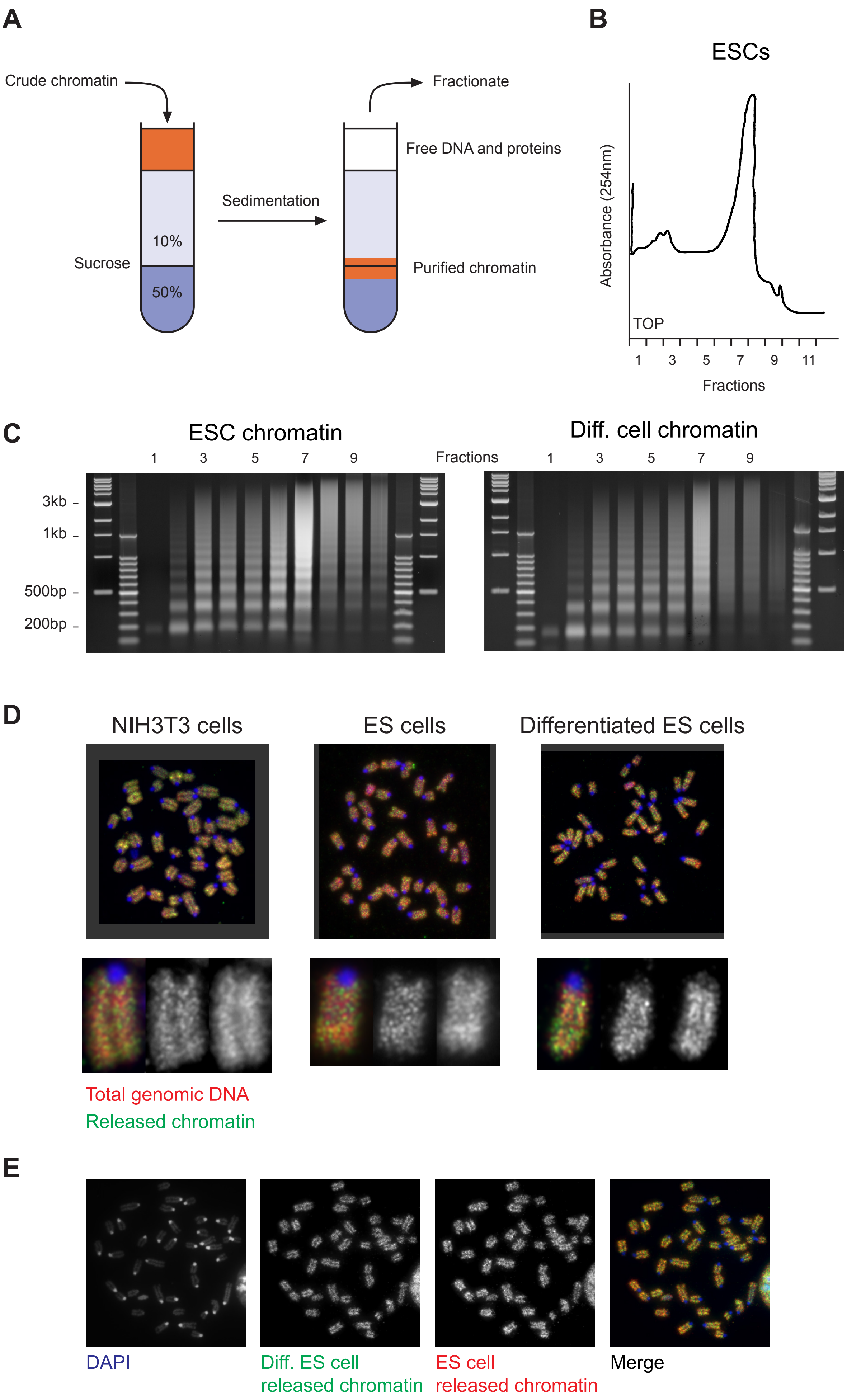


Gilbert Supplementary Figure 1

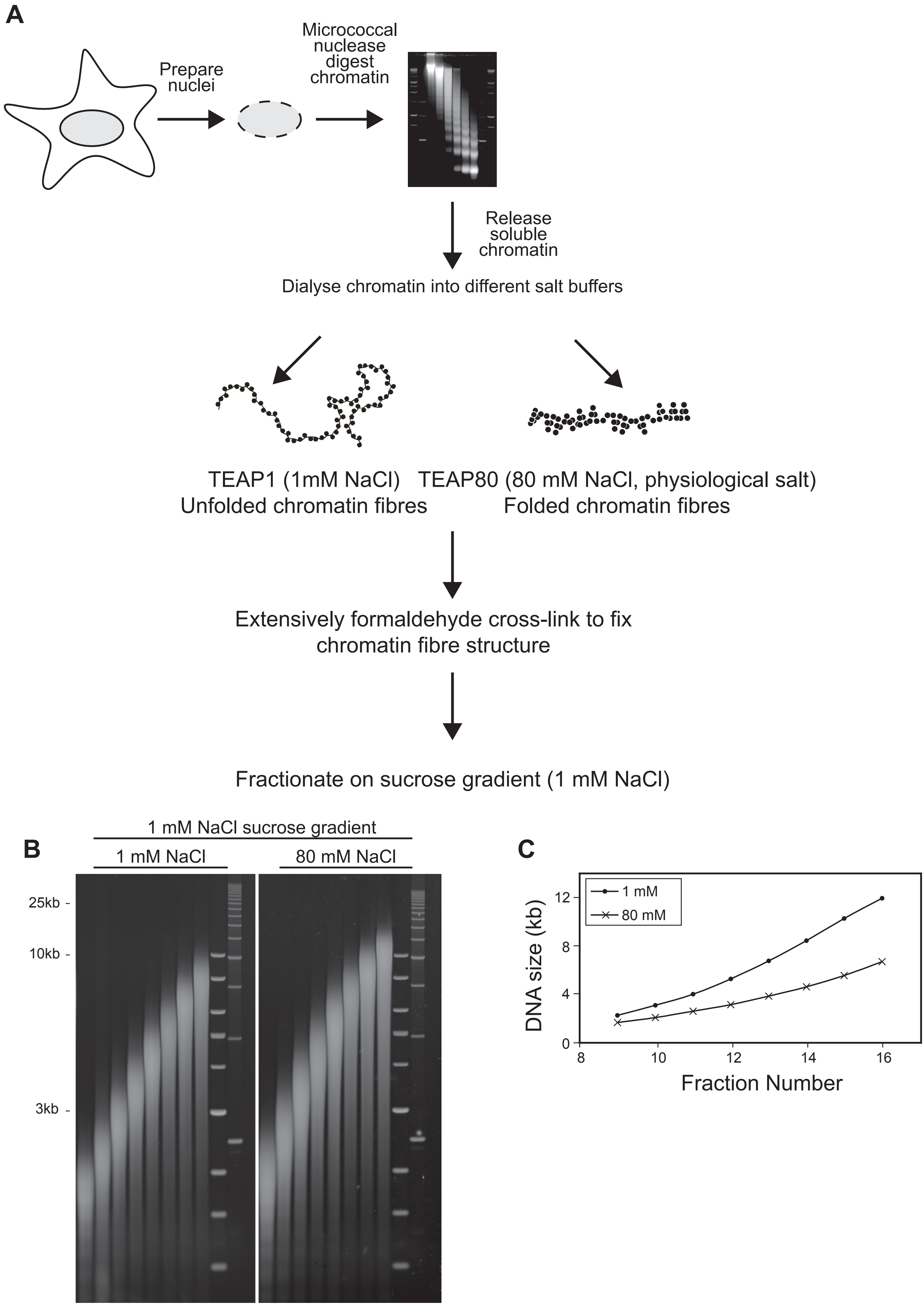


Gilbert Supplementary Figure 2

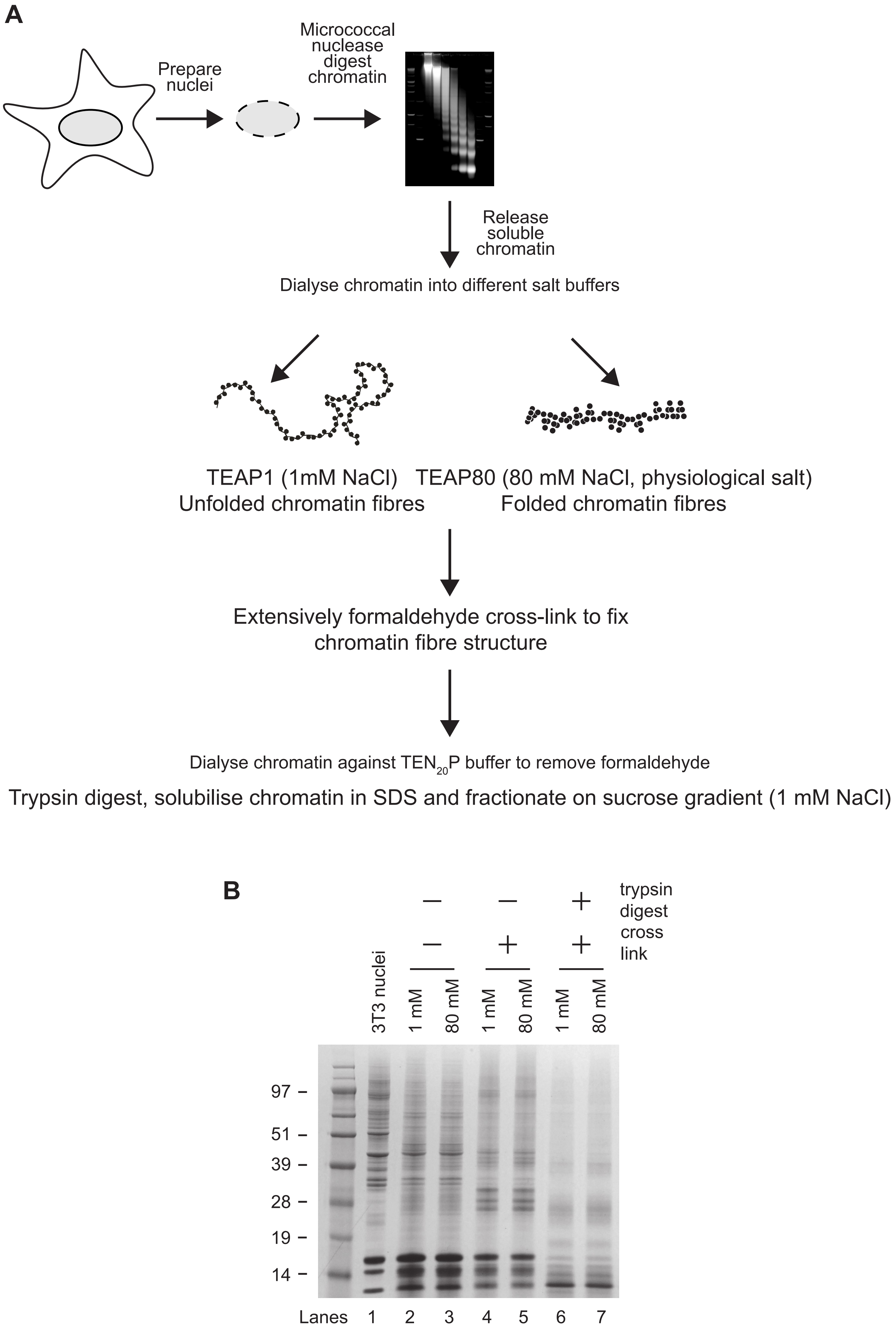


Gilbert. Supplementary Figure 3

C

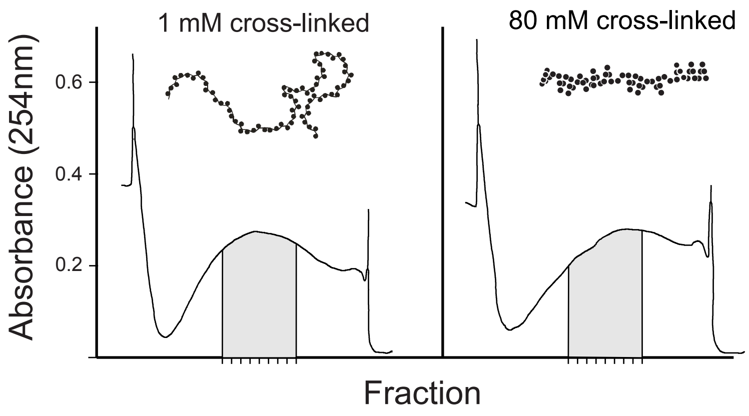


D

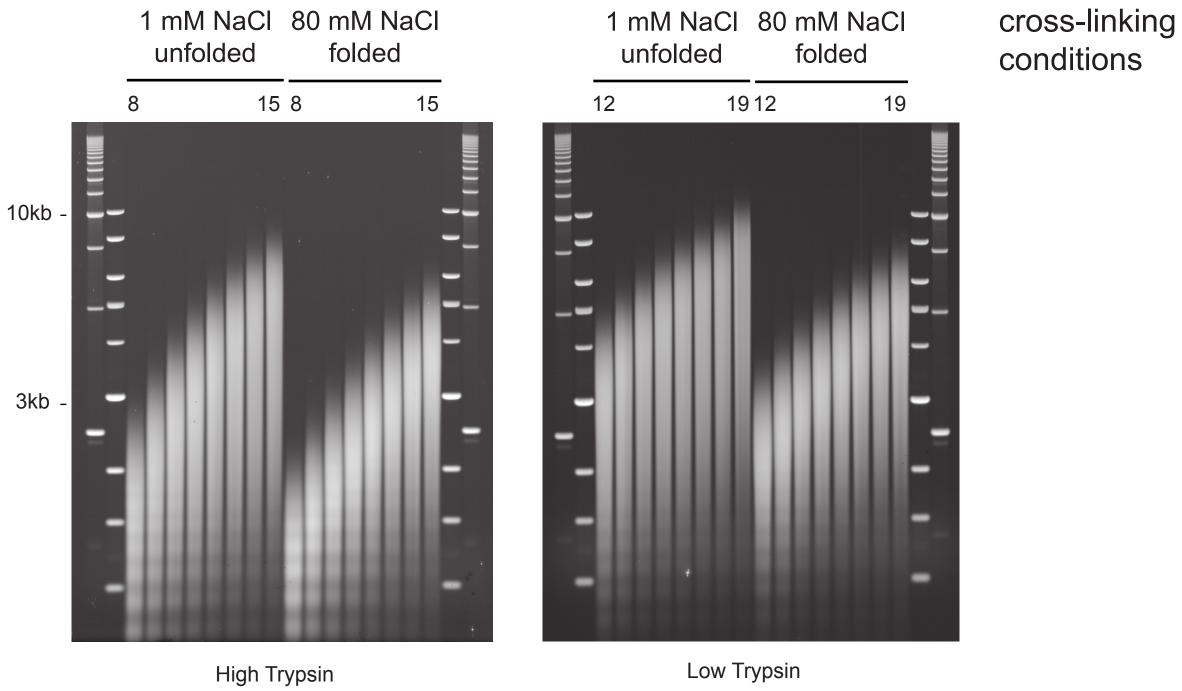


E

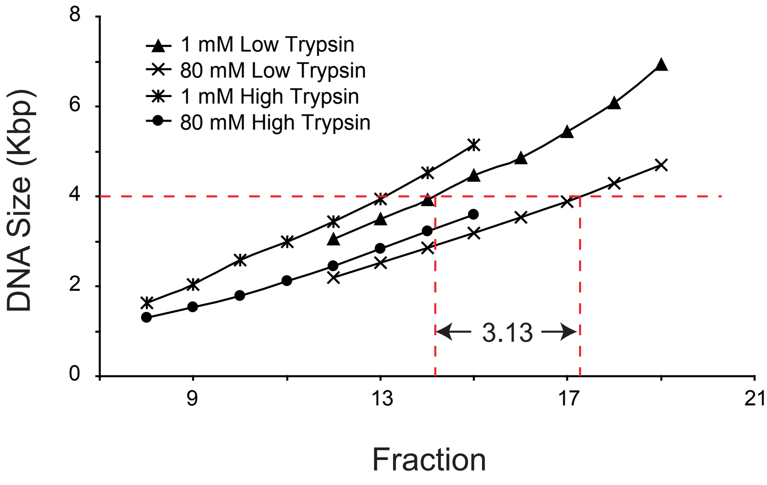


Gilbert Supplementary Figure 3

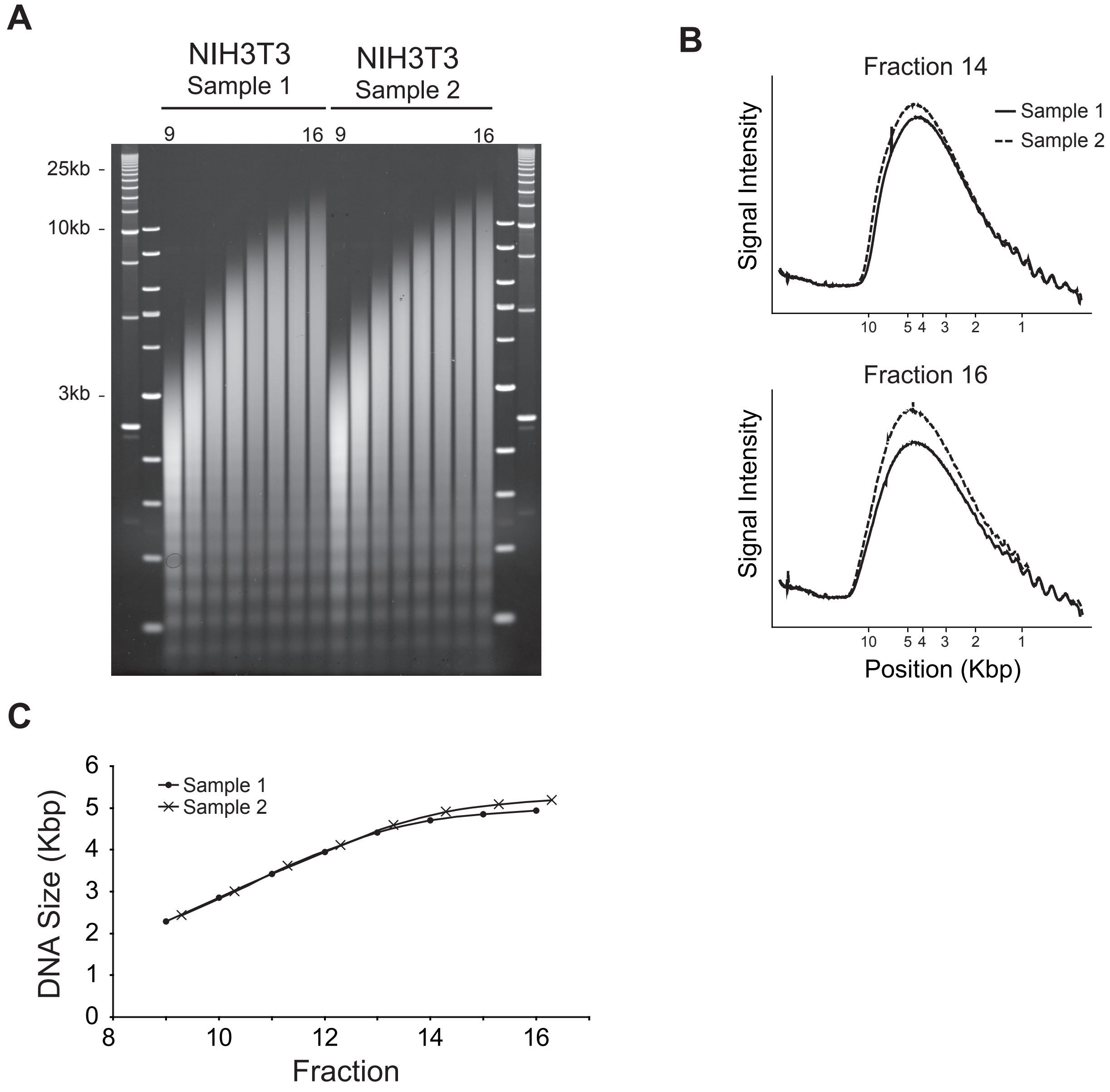


Gilbert Supplementary Figure 4

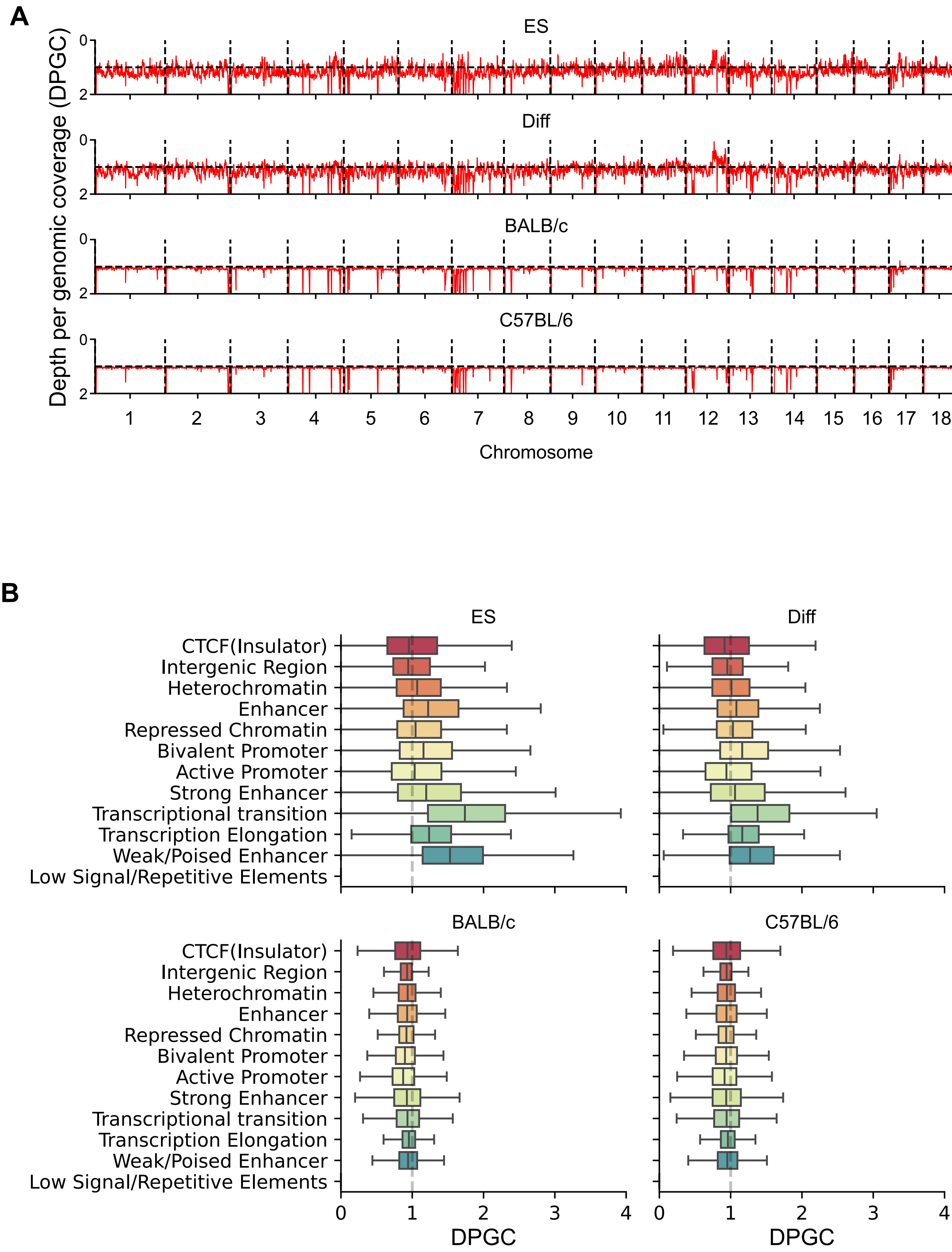


Gilbert Supplementary Figure 5
